# Comparative analysis of sequencing technologies platforms for single-cell transcriptomics

**DOI:** 10.1101/463117

**Authors:** Kedar Nath Natarajan, Zhichao Miao, Miaomiao Jiang, Xiaoyun Huang, Hongpo Zhou, Jiarui Xie, Chunqing Wang, Shishang Qin, Zhikun Zhao, Liang Wu, Bo Li, Yong Hou, Shiping Liu, Sarah A. Teichmann

## Abstract

All single-cell RNA-seq protocols and technologies require library preparation prior to sequencing on a platform such as Illumina. Here, we present the first report to utilize the BGISEQ-500 platform for scRNA-seq, and compare the sensitivity and accuracy to Illumina sequencing. We generate a scRNA-seq resource of 468 unique single-cells and 1,297 matched single cDNA samples, performing SMARTer and Smart-seq2 protocols on mESCs and K562 cells with RNA spike-ins. We sequence these libraries on both BGISEQ-500 and Illumina HiSeq platforms using single- and paired-end reads. The two platforms have comparable sensitivity and accuracy in terms of quantification of gene expression, and low technical variability. Our study provides a standardised scRNA-seq resource to benchmark new scRNA-seq library preparation protocols and sequencing platforms.

## Background

Single cell RNA-seq (scRNA-seq) has become the established approach to dissect cellular heterogeneity, unravel cell states and identify subpopulation structures across different cell types [1–4]. The different scRNA-seq methods and technologies have been benchmarked using synthetic RNA spike-ins [5–7]. However, to date, most scRNA-seq methods require cDNA libraries to be compatible with Illumina sequencing platform. BGISEQ-500 provides an alternative short-read sequencing technology, which is based on combinatorial probe-anchor synthesis (cPAS) to form DNA nanoball nanoarrays for stepwise sequencing using polymerase [13]. The Illumina system uses stepwise sequencing by polymerase on DNA microarrays prepared by bridge PCR. Here, we assess the suitability of BGISEQ-500 sequencing platform for scRNA-seq, and compare with matched data obtained from the Illumina HiSeq platform.

We perform two different scRNA-seq methods (SMARTer and Smart-seq2) on mouse embryonic stem cells and human K562 cells [8,9]. We utilize both External RNA Controls Consortium (ERCCs) and Spike-in RNA Variants (SIRVs) that span 92 synthetic RNA species and 69 artificial transcripts, of varying lengths, concentrations, GC contents, isoforms and abundance levels. We benchmark and compare the performance metrics (*sensitivity* and *accuracy*) for the different cells, protocols and technologies across both sequencing platforms.

As in our previous framework [5], we define ‘sensitivity’ (or lower detection limit) as the minimum number of input RNA spike-in molecules that are detected as expressed within a single-cell. ‘Accuracy’ refers to the correlation between the estimated abundances of input RNA spike-ins and the ground truth, *i.e.* input molecules added to the single-cell reaction (Fig. S1A-B). The BGISEQ-500 platform has been previously applied to detection of small noncoding RNAs [10], human genome re-sequencing [11] and palaeogenomic ancient DNA sequencing [12], but not to scRNA-seq.

In this study, we perform the first systematic scRNAseq comparison across the two sequencing platforms, using 1,297 matched single cDNA samples from 468 unique single-cells across different scRNA-seq protocols. We compare and assess whether alternative library preparation and sequencing platform can be utilized for the same scRNA-seq samples with comparable accuracy, sensitivity and robustness as the current state-of-art Illumina HiSeq platform. The generated scRNA-seq datasets also provide a large data resource for comparison of new protocols and computational methods.

## Results

We performed two scRNA-seq methods (SMARTer and Smart-seq2) in parallel on 288 single-mESCs using both ERCCs and SIRVs spike-ins on the microfluidics-based Fluidigm C1-system [8,9]. The Smart-seq2 method was performed in replicates (SM2 replicate 1 and 2). The single-cell lysis, reverse transcription and pre-amplification for all methods was done within the C1-system. We used the single-cell cDNA from each protocol to generate 576 single-cell sequencing libraries for Illumina HiSeq2500 (Illumina) and BGISEQ-500 (Fig. 1A; Methods). This allowed us to sample the same *‘matched’* single-cells (288 cells) across two sequencing platforms (Fig. 1A).

**Figure 1:**
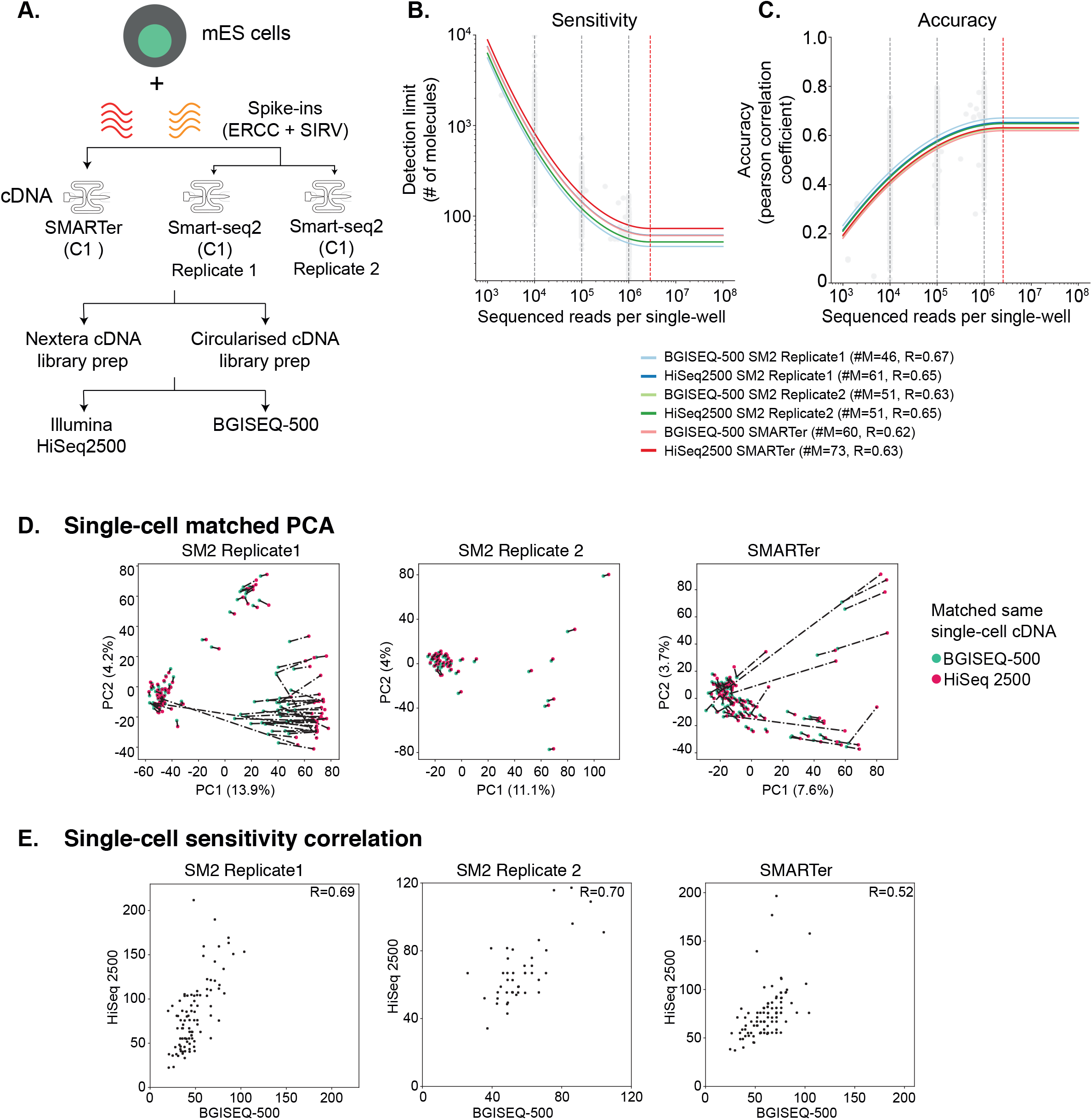
**(A)** Schematic overview of the mESC scRNA-seq experiment and sequencing. **(B)** Single-cell detection limit (Sensitivity) of mESC cells, downsampled across two orders of magnitude from SMARTer and two Smart-seq2 replicates (633 samples). The single-cell sensitivities are largely similar between different library preparation across scRNA-seq protocols. **(C)** Single-cell accuracy of mESC cells, downsampled across two orders of magnitude for SMARTer and two Smart-seq2 replicates (633 samples). The grey dotted lines indicate downsampling at different read depths per cell, while red line indicates saturation per cell. **(D)** PCA for matched single-cell cDNA samples performed using SMARTer and two replicates of Smart-seq2 and sequenced across both sequencing platforms. Red and green colored circles indicate sequencing of matched cDNA on Illumina HiSeq2500 and BGISEQ-500 respectively. The dotted lines represent distance i.e. measure of similarity across sequencing platforms. **(E)** Single-cell correlations for each scRNA-seq protocol and across sequencing platforms. The correlations (*R*=0.52~0.70) are comparable between sequencing platforms.

For each single-cell, we performed filtering, quality control and converted aligned reads to normalized transcript per million (TPM) units (Methods). We calculated the sensitivity and accuracy for both spike-ins across matched single-cells sequenced on both sequencing platforms, as devised in our past computational framework [5] (Fig. S1C-D). Globally, the single-cell accuracy was similar across sequencing platforms, irrespective of scRNA-seq protocol (Pearson correlation coefficient *R*=0.66-0.70, Fig. S1C, E). The sensitivity (i.e. detection limit) was comparable, ranging from 21 to 47 molecules (number of RNA molecules *#M*=21-47; Fig. S1D, F) between sequencing platforms, while SM2-seq replicate 1 could detect down to 12 RNA molecules within single cells (Fig. S1F).

Since sensitivity can be dependent on sequencing depth [5], we compared whether the total number of sequenced reads varied between platforms. Not surprisingly, the BGI-cells had much higher sequencing depth, which accounts for slightly increased sensitivity (Fig. S1G). At the respective sequencing depths, the number of genes expressed across single mESCs were similar across scRNA-seq protocol and sequencing platform (Fig. S1H). Interestingly, the SM2 replicate 1 could be split into two robust subpopulations across both sequencing platforms, differing in number of genes expressed and expression of key pluripotency markers (Fig. S1H). The truly pluripotent subpopulation had increased global expression and of stem cell markers (Illumina>5000 and BGI>7500 genes), while the differentiated-like subpopulation expressed fewer genes and stem cell markers (Fig. S1HI) [14]. The statistics for each matched single-cell are summarised in Table S1.

To account for biases due to sequencing depth and technical variability, we downsampled total reads across two orders of magnitude (raw reads to 10^6^, 10^5^, 10^4^ total reads). We re-computed sensitivity and accuracy using a linear model that returns individual corrected performance parameters on the data. After accounting for depth, both sensitivity and accuracy were highly similar across single-cells between scRNA-seq protocols and sequencing platforms (Fig. 1BC). The detection limit was consistent between scRNA-seq protocols and sequencing platforms (*#M*=46- 73), with the sensitivity reaching saturation around ~3 million reads (red dashed line; Fig. 1B). The accuracy was also consistent (*R*=0.62~0.67) between platforms, reaching saturation at ~2.5 million reads (red dashed line; Fig. 1C).

Since the same single cells were sequenced across both platforms, we next assessed the matched cell-similarity by comparing total expression (Genes+spike-ins), and for spike-ins alone, by random downsampling to 1 million reads per single-cell. First, we performed Principal component analysis (PCA) using total gene expression, plotting each matched cell by two points representing the sequencing platforms, where the distance between them is a measure of gene expression similarity (dashed line; Fig 1D). The PCA captured strong similarity between matched cells (short distance) for most of the single-cells across both platforms, with low PC1 (8-14%) and PC2 (4%) contribution.

To control for technical noise and selective bias from genes, we also performed PCA using both spike-ins (without genes) on matched cells. The matched cells in resulting PCA were more uniformly place with short distance and low PC1 (~7%) and PC2 (~5-6%) variation across both platforms (Fig. S2A). For SM2 replicate 1, we overlaid the PCA with number of gene expressed and could re-confirm that subpopulations were biological and not driven by technical variation (Fig. S1I, S2B). To assess the performance metrics between the two sequencing platforms, we correlated the single-cell sensitivities for each matched cell both before and after downsampling. Without any downsampling, we expected and observed that the single-cell sensitivity correlations were low, owing to dependence on sequencing depth (Fig S2C). However upon downsampling to 1 million reads, the correlations were significantly improved and were more consistent across protocols and sequencing platforms (*R*=0.52~0.70). In summary, the comparison of different performance metrics across matched single-cells using different protocols was highly comparable and similar between Illumina HiSeq2000 and BGISEQ-500 sequencing platforms.

We repeated our benchmarking comparison on plate-based Smart-seq2 using 82 mESCs and 98 human K562 cells, containing ERCC spike-ins only. From both mESCs and K562s, we generated 600 sequencing libraries for BGISEQ-500 using both single- and paired-end sequencing and *matched* 121 sequencing libraries for HiSeq 4000 paired-end sequencing (721 sequencing libraries in total) (Fig. 2A; Table S1-S5). We generated 3-20 million reads for each single across both sequencing platforms. To account for sequencing depth, we randomly downsampled to 1 million reads per cell and calculated performance metrics for each single-cell across sequencing platforms. The mESCs and K562s have dramatically variable amount of cellular RNA/cell, however we could robustly detected >8500 genes and spike-ins in each cell type, with over 90% commonly detected between both platforms (Fig. S3A; Table S2). Owing to differential RNAs/cell, the K562 had a increased sensitivity across both sequencing platforms and we could consistently detect genes and spike-ins represented by just a few molecules (#M=3-4). This compares to a somewhat worse but consistent mESCs detection limit (#M=42~49). The accuracy for both cell types was quite high and consistent (*R*=~0.85~0.88), indicating similar performance metrics across sequencing platforms (Fig. 2B and S3B-C). The statistics for each matched K562 and mES cell across both platforms are provided in Table S2.

**Figure 2:**
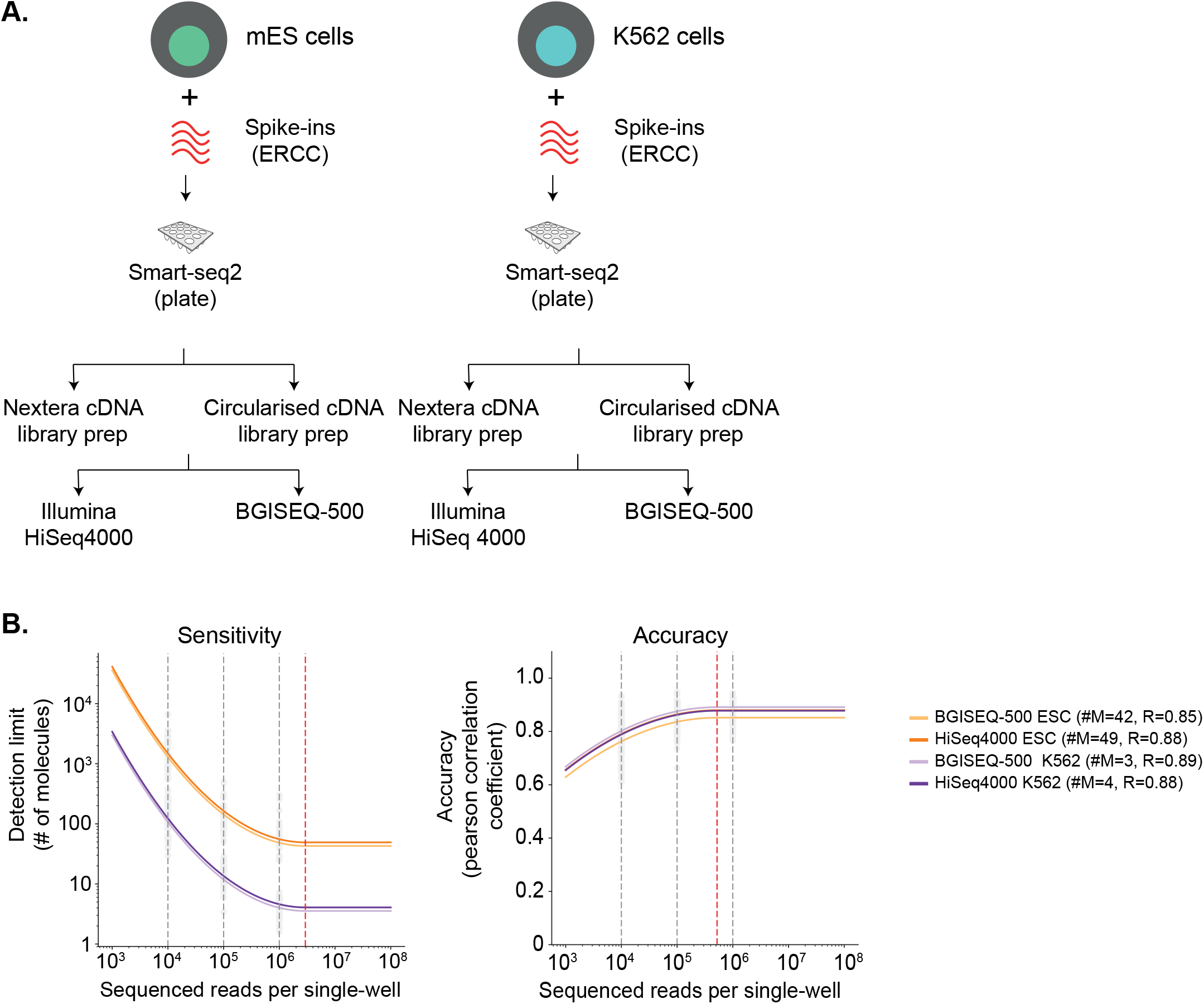
**(A)** Schematic overview of the mESC and K562 scRNA-seq experiment using plate-based Smart-seq2 protocol and sequencing. **(B)** The sensitivity of mESCs and K562s cells downsampled across two orders of magnitude (633 samples). The sensitivity is critically dependent on sequencing depth. Therefore, more deeply sequenced K562 are more sensitive than mESCs. Between the sequencing platforms, sensitivity is highly similar **(C)** The accuracy of mESCs and K562s downsampled across two orders of magnitude (633 samples). The accuracy is marginally dependent on sequencing depth. Therefore, accuracy is largely similar and comparable between mESCs and K562s, irrespective of sequencing platform. The grey dotted lines indicate downsampling at different read depths per cell, while red line indicates saturation per cell.

Since we generated mESC data from different scRNA protocols, technologies and sequencing platforms containing ERCC spike-ins in two different batches, we collectively assessed the mESCs performance metrics for both platforms. We downsampling the raw reads and observed that both sensitivity and accuracy were comparable between both platforms (Fig. S3DE). The sensitivity and accuracy were saturated at ~2 million and ~200,000 reads respectively (Fig. S3A-B). We also observed similar detected genes detected between both platforms (Fig. S3F, Table S2). In summary, our analysis demonstrates similar and robust performance metrics between BGISEQ-500 and Illumina platforms for scRNA-seq.

Our dataset spans 468 unique single cells of two different cell types (mESCs, K562s), two scRNA-seq protocols (SMARTer, Smart-seq2), two technologies (Fluidigm C1, plate-based) and matched 1,297 libraries across Illumina and BGISEQ-500 sequencing platform (Table S1-S4). In addition to paired-end sequencing data across BGISEQ-500 and Illumina platform, we also generated matched mESCs and K562 scRNA-seq data using 100bp single-end (SE), generating a huge benchmarking data with over 750 GB of raw single-cell data. Comparing the BGISEQ-500 SE and PE data, the average sequencing depth per cell was ~9.6 and ~8.7 million reads respectively (Table S3 and Table S4). Both SE and PE datasets detected >9500 genes for K562s and >9000 genes for mESCs. The accuracies for K562 and mESCs cells across both SE and PE reads was *R*=~0.70-0.85, and the sensitivities for K562 and mESCs cells were #M=4~25 across both SE and PE reads. Comparing the raw reads across both mESCs and K562, we also observed higher number alternative splicing events at single-cell level across all sequencing runs (Fig. S3G).

In summary, our datasets and comparative performance assessment offers a large standardized resource to the community to further investigate potential technical biases including GC content, isoform quantification, impact of read-lengths across different scRNA-seq protocols, technologies and sequencing platforms. The matched 1,297 single-cell datasets and annotations would serve as an ideal starting point for benchmarking and comparison of new protocols and computational methods for the scientific community.

## Discussion

The rapid developments in single-cell genomics are transforming our understanding of biological systems by capturing underlying gene expression variability to identify cell types, states and transitions across cell populations. Single-cell transcriptomic profiling is a multi-step sampling procedure, where the first major step involves cell lysis, RNA capture, reverse-transcription of RNA, preamplification of cDNA generation. The next major step requires single-cell cDNA to be converted into a sequencing compatible library, followed by sequencing. There are several scRNA-seq protocols that utilize different chemistries, platforms and technologies to address the first critical step of converting RNA into cDNA. The technical variation, performance metrics (sensitivity, accuracy) and reproducibility for the first critical step have been recently evaluated and benchmarked using synthetic RNA spike-in molecules [5–7]. However, all the scRNA-seq protocols and technologies require libraries to be compatible for sequencing on Illumina platform.

Our study is the first to utilize BGISEQ-500 platform for scRNA-seq. Our comprehensive benchmarking of performance metrics utilises two scRNA-seq protocols (SMARTer and Smart-seq2), multiple spike-ins (ERCC alone, ERCC+SIRV), two different cell lines (mESCs, K562s), two technologies (Fluidigm C1, plate-based) across Illumina HiSeq and BGISEQ-500 platform. Utilizing 468 single K562 and mES cells and matched 1,297 single-cell libraries, we observe BGISEQ-500 to be highly comparable in sensitivity, accuracy and reproducibility to Illumina platform, while being considerably more cost effective.

From our mESC scRNA-seq dataset, we could further distinguish technical artifacts (sequencing depth) from biological variation (subpopulations) across both sequencing platforms. We observe higher alternative splicing events in K562s compared to mESCs across both sequencing platforms. Our data strongly supports the notion that minimal variability is introduced during library preparation and sequencing for both Illumina and BGISEQ-500 platforms. In combination with our previous framework [5], we believe that variability between the steps of scRNA-seq protocols is largest during the RNA to cDNA step. Given the similar single-cell characteristics (number of genes, expression range, subpopulation etc.) and performance metrics across different library preparation and sequencing platforms in our datasets, BGISEQ-500 library preparation and sequencing is robust and comparable to Illumina HiSeq platforms for single-cell applications.

We observe minimal variability in cDNA processing across different library preparation and sequencing platforms. In the current study, we did not perform scRNA-seq protocols with Unique Molecular Identifiers (UMIs) that account for PCR amplification biases. Given that UMIs primarily address biases during the RNA-to-cDNA stage (and to cDNA amplification), this would not impact our assessment of sequencing platforms. The UMI protocols could easily be extended to be compatible with sequencing on BGISEQ-500 platform. Our large resource for benchmarking scRNA-seq data suggests that the BGISEQ-500 platform is suitable for plate-based (microwell or nanowell), droplet and microfluidics technologies.

In addition to benchmarking, we provide a large comprehensive multi cell type, protocol and platform scRNA-seq dataset spanning 468 cells and 1,297 libraries to the community. Given the large research initiatives profiling transcriptomes of single cells in mouse [3,15] and Human, such as the Human Cell Atlas [16], achieving high quality and cost effective methods is paramount. Our standardized resource can be utilized for investigating technical biases and for benchmarking scRNA-seq protocols and computational methods.

## Declarations

### Data availability

The datasets generated and analysed during the current study are deposited at ArrayExpress (E-MTAB-7239), BioProject (PRJNA430491) and SRA (SRP132313).

## Acknowledgements

The authors thank the Teichmann lab for helpful discussions and comments on the manuscript.

## Authors’ contributions

KNN, XH, SL and SAT designed and supervised the project. KNN, HZ, JX and CW performed the experiments with help from SQ, ZZ, LW, BL, and YH. KNN and ZM performed the bioinformatics analysis. KNN, ZM, MJ, XH and SAT wrote the manuscript. All authors reviewed and approved the manuscript.

---

**Supplementary Figure 1:**
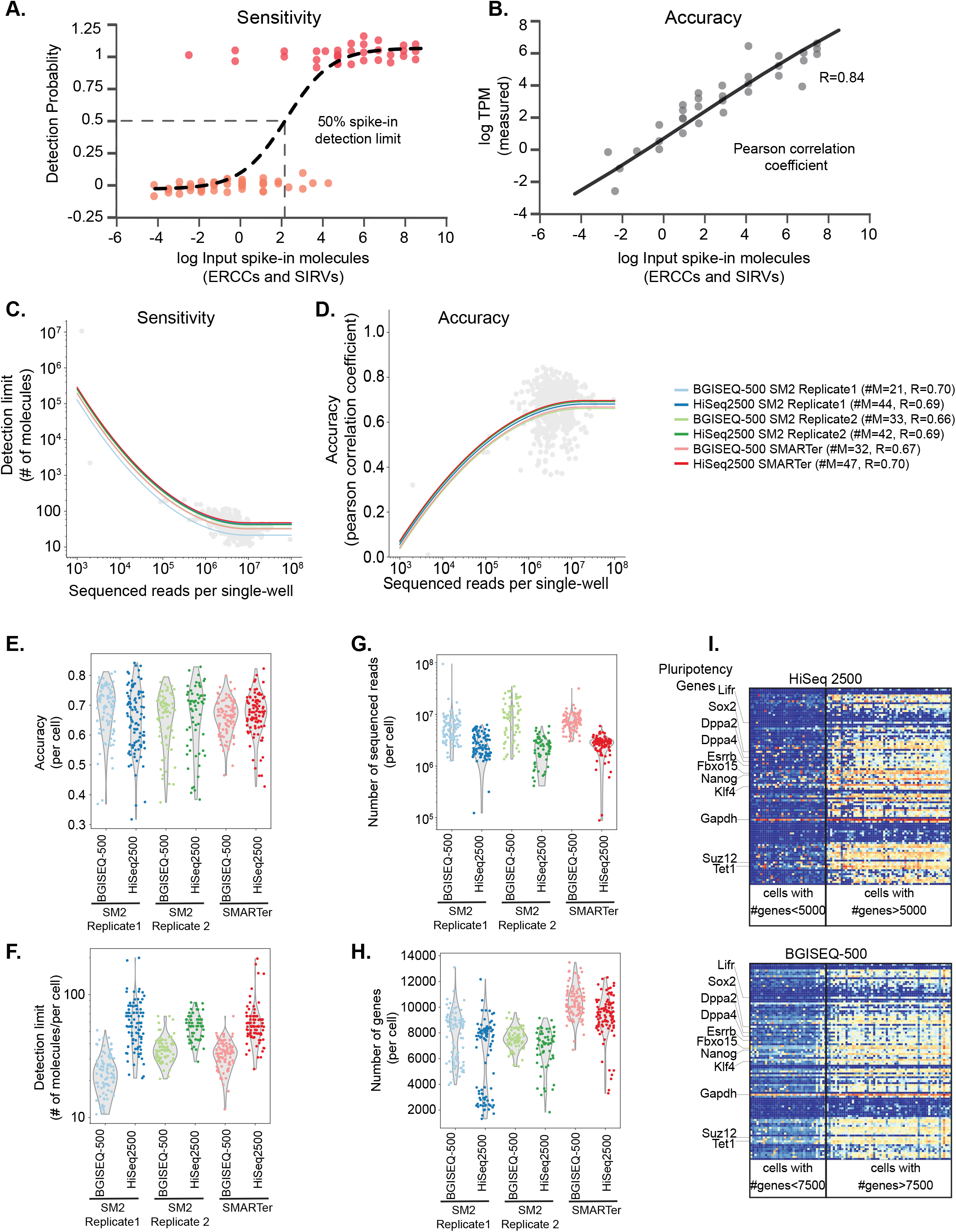
**(A)** Schematic description of performance metrics for comparing scRNA-seq protocols and sequencing platforms. Sensitivity is described as the minimum number of input molecules (spike-in), where the detection probability is 50%. **(B)** Accuracy is calculated as the Pearson correlation between estimated expression (TPMs) and input spike-in concentrations (ground-truth). **(C&D)** Single-cell accuracy and sensitivity of mESC cells (without downsampling) for SMARTer and two Smart-seq2 replicates (633 samples) across both sequencing platforms. **(E-H)** Single-cell accuracy, sensitivity, sequenced reads and genes detected between scRNA-seq protocols and sequencing platforms represented in violin plots. Two subpopulations are observed in Smart-seq2 replicate 2 based on genes detected across both sequencing platforms. **(I)** Heatmap with expression of pluripotency markers for Smart-seq2 replicate 2 cells across both HiSeq2500 and BGISEQ-500 cells. The cutoff of 5000 (HiSeq2500) and 7500 genes (BGISEQ-500) was visually selected based on Fig S1H. The subpopulation expressing higher number of genes is truly pluripotent, while the other subpopulation seems to be on differentiation path.

**Supplementary Figure 2:**
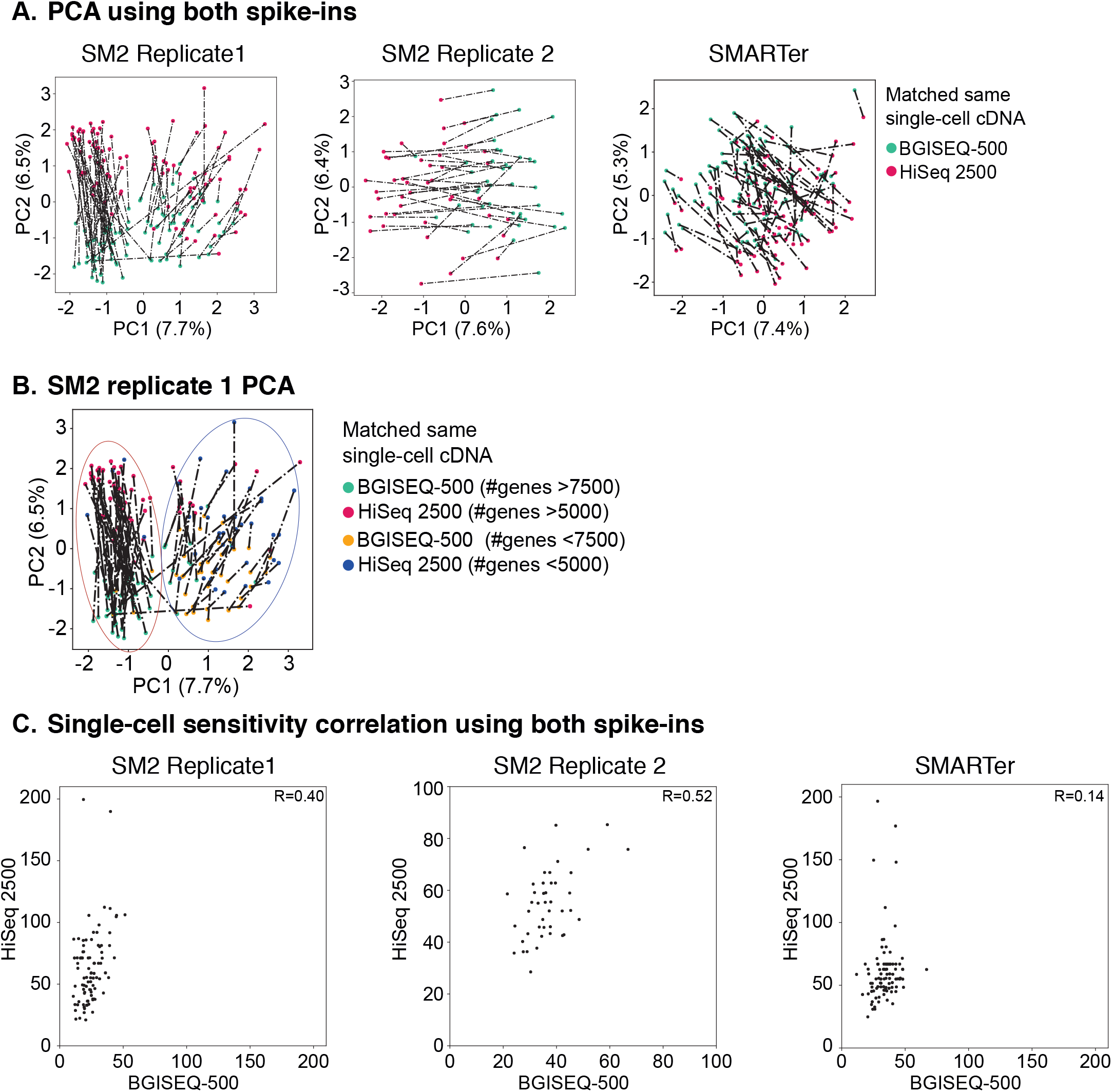
**(A)** PCA using only spike-in RNAs (ERCCs and SIRVs) for matched single-cell cDNA samples performed using SMARTer and two replicates of Smart-seq2 and sequenced across both sequencing platforms. Red and green colored circles indicate sequencing of matched cDNA on Illumina HiSeq2500 and BGISEQ-500 respectively. The dotted lines represent distance i.e. measure of similarity across sequencing platforms. **(B)** PCA of Smart-seq2 replicate 2 re-colored by subpopulations with different number of genes detected. The cutoff of 5000 (HiSeq2500) and 7500 genes (BGISEQ-500) was visually selected based on Fig S1H. **(C)** Single-cell correlations using only spike-in RNAs (ERCCs and SIRVs) for each scRNA-seq protocol and between sequencing platforms. The correlations (*R*=0.14~0.52) are quite poor owing to critical dependence of sensitivity on sequencing depth.

**Supplementary Figure 3:**
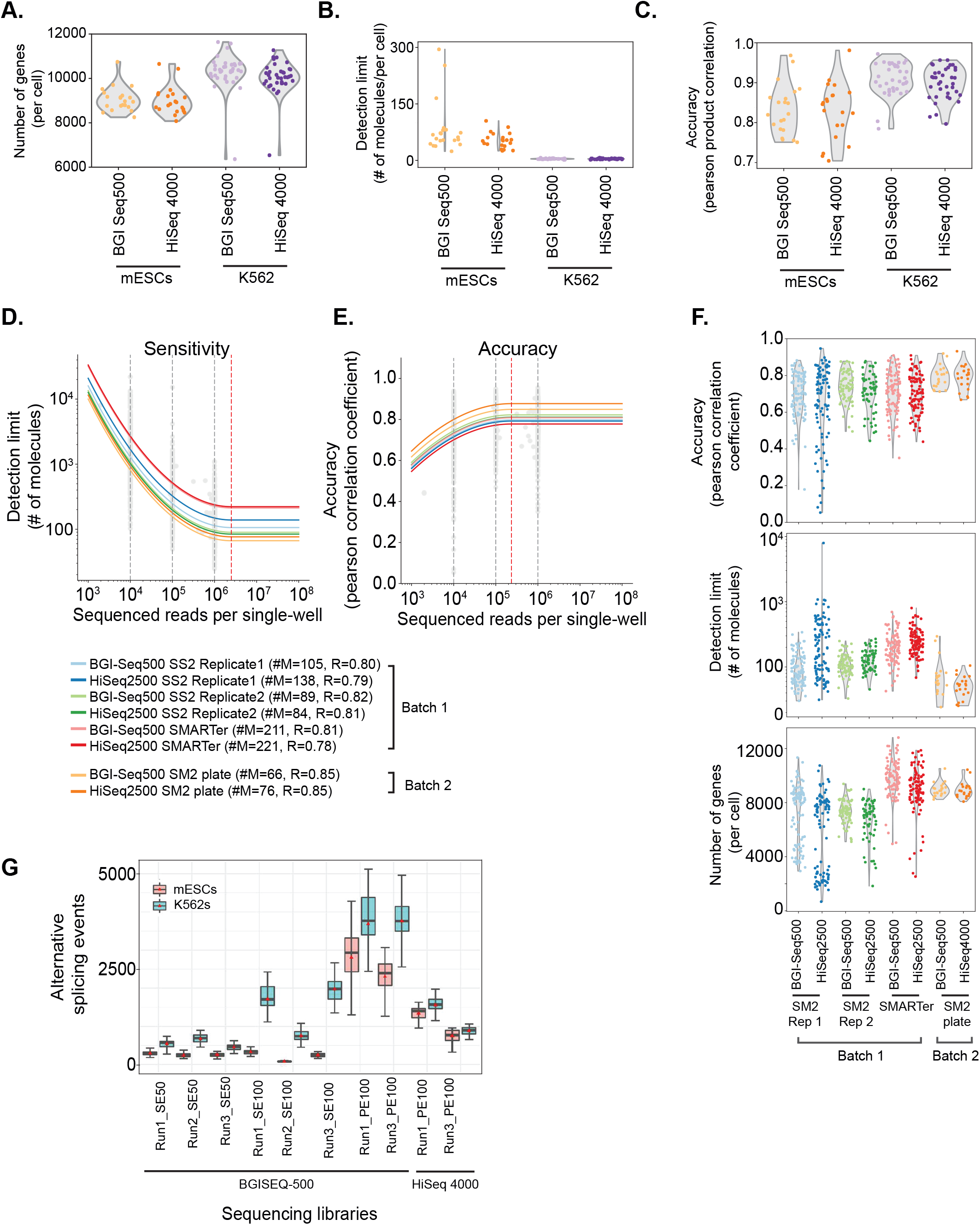
**(A-C)** Number of genes detected **(A)**, sensitivity **(B)** and accuracy **(C)** for mESCs and K562s between both sequencing platforms represented in violin plots. Higher sequencing depth of K562s accounts for higher gene detection, higher sensitivity and more accuracy. **(D-F)** Comparison of all mESC data from different scRNA-seq protocols. The sensitivity **(D)** and accuracy **(E)** single-cell comparison between protocols and sequencing platforms after downsampling. The grey dotted lines indicate downsampling (10^4^, 10^5^, 10^6^ reads), while red line indicates saturation per cell. **(F)** Visualisation of single-cell accuracy, detection limit and genes detected (without downsampling) in violin plots. The accuracy is largely similar for all protocols and sequencing platforms, while the sensitivity and genes detected vary with sequencing depth. **(G)** Box plots describing number of alternative splicing events across mESCs and K562s using raw reads from all sequencing runs. The SE and PE describe single- and paired-end sequencing with respective read lengths across both sequencing platforms.

**Supplementary Table 1:** Single-cell library statistics computed from raw sequencing reads for mESCs and K562s performed using SMARTer and Smart-seq2 protocols and sequenced across HiSeq2500 and BGISEQ-500 platforms.

**Supplementary Table 2:** Single-cell library statistics computed from randomly downsampled 1 million sequencing reads for mESCs and K562s from both scRNA-seq protocols and across HiSeq2500 and BGISEQ-500 sequencing platforms.

**Supplementary Table 3:** Single-cell library statistics computed from for all mESCs and K562 replicate libraries sequenced on BGISEQ-500 platform using single-end (SE) reads

**Supplementary Table 4:** Single-cell library statistics computed from for all mESCs and K562 replicate libraries sequenced on BGISEQ-500 platform using Paired-end (SE) reads.

## References

1. Tirosh I, Izar B, Prakadan SM, Wadsworth MH 2nd, Treacy D, Trombetta JJ, et al. Dissecting the multicellular ecosystem of metastatic melanoma by single-cell RNA-seq. Science. 2016;352:189–96.

2. Lönnberg T, Svensson V, James KR, Fernandez-Ruiz D, Sebina I, Montandon R, et al. Single-cell RNA-seq and computational analysis using temporal mixture modelling resolves Th1/Tfh fate bifurcation in malaria. Sci Immunol [Internet]. 2017;2. Available from: http://dx.doi.org/10.1126/sciimmunol.aal2192

3. Han X, Wang R, Zhou Y, Fei L, Sun H, Lai S, et al. Mapping the Mouse Cell Atlas by Microwell-Seq. Cell. 2018;173:1307.

4. Natarajan KN, Teichmann SA, Kolodziejczyk AA. Single cell transcriptomics of pluripotent stem cells: reprogramming and differentiation. Curr Opin Genet Dev. 2017;46:66–76.

5. Svensson V, Natarajan KN, Ly L-H, Miragaia RJ, Labalette C, Macaulay IC, et al. Power analysis of single-cell RNA-sequencing experiments. Nat Methods. 2017;14:381–7.

6. Ziegenhain C, Vieth B, Parekh S, Reinius B, Guillaumet-Adkins A, Smets M, et al. Comparative Analysis of Single-Cell RNA Sequencing Methods. Mol Cell. 2017;65:631–43.e4.

7. Wu AR, Neff NF, Kalisky T, Dalerba P, Treutlein B, Rothenberg ME, et al. Quantitative assessment of single-cell RNA-sequencing methods. Nat Methods. 2014;11:41–6.

8. Picelli S, Faridani OR, Björklund AK, Winberg G, Sagasser S, Sandberg R. Full-length RNA-seq from single cells using Smart-seq2. Nat Protoc. 2014;9:171–81.

9. Ramsköld D, Luo S, Wang Y-C, Li R, Deng Q, Faridani OR, et al. Full-length mRNA-Seq from single-cell levels of RNA and individual circulating tumor cells. Nat Biotechnol. 2012;30:777–82.

10. Fehlmann T, Reinheimer S, Geng C, Su X, Drmanac S, Alexeev A, et al. cPAS-based sequencing on the BGISEQ-500 to explore small non-coding RNAs. Clin Epigenetics. 2016;8:123.

11. Huang J, Liang X, Xuan Y, Geng C, Li Y, Lu H, et al. A reference human genome dataset of the BGISEQ-500 sequencer. Gigascience. 2017;6:1–9.

12. Mak SST, Gopalakrishnan S, Carøe C, Geng C, Liu S, Sinding M-HS, et al. Comparative performance of the BGISEQ-500 vs Illumina HiSeq2500 sequencing platforms for palaeogenomic sequencing. Gigascience. 2017;6:1–13.

13. Drmanac R, Sparks AB, Callow MJ, Halpern AL, Burns NL, Kermani BG, et al. Human genome sequencing using unchained base reads on self-assembling DNA nanoarrays. Science. 2010;327:78–81.

14. Kolodziejczyk AA, Kim JK, Tsang JCH, Ilicic T, Henriksson J, Natarajan KN, et al. Single Cell RNA-Sequencing of Pluripotent States Unlocks Modular Transcriptional Variation. Cell Stem Cell. 2015;17:471–85.

15. The Tabula Muris Consortium, Quake SR, Wyss-Coray T, Darmanis S. Single-cell transcriptomic characterization of 20 organs and tissues from individual mice creates a Tabula Muris [Internet]. 2017. Available from: http://dx.doi.org/10.1101/237446

16. Rozenblatt-Rosen O, Stubbington MJT, Regev A, Teichmann SA. The Human Cell Atlas: from vision to reality. Nature. 2017;550:451–3.

